# A parenchymal niche regulates pluripotent stem cell function in planarians

**DOI:** 10.1101/2025.08.01.668211

**Authors:** Skylar E. Settles, Kathleen E. Miller, Rachel H. Roberts-Galbraith

## Abstract

Harnessing the extraordinary potential of stem cells requires exquisite control *in vitro*, *in vivo*, and in the context of potential therapies. Adult stem cells are commonly housed within a microenvironment, or “niche,” that supplies signals to regulate stem cell function. It has been difficult to determine whether pluripotent stem cells have a similar endogenous niche due to the transient nature of the cell type in most animal species. We sought to establish the nature of a pluripotent stem cell niche using planarian flatworms, animals that maintain pluripotent stem cells throughout adulthood and use them for whole-body regeneration. We characterized the cellular microenvironment of planarian stem cells in detail, finding that each stem cell is surrounded by a diverse mixture of other stem cells and differentiated cells. We identified phagocytic cells marked by *CTSL2* as the differentiated cell type most frequently found next to stem cells, in both homeostasis and regeneration. *CTSL2^+^* cells, which we named abraçada cells, wrap around stem cells, and then neighbor progenitors less often coincident with their specialization. We then tested whether abraçada cells constitute a functional niche that regulates stem cells. We identified two innexin-encoding genes expressed in abraçada cells that are critical for pluripotent stem cell function and planarian regeneration. Together, our results reveal for the first time a local, endogenous niche that actively regulates adult pluripotent stem cells.

## Introduction

In most animal species, adult stem cells are uni- or multipotent and reside within a tissue-specific, specialized microenvironment called a “niche.” A niche regulates stem cell behaviors including proliferation, quiescence, and differentiation [1]. When differentiated cells of a niche become dysfunctional, stem cells become depleted or abnormal. For example, in the *Drosophila* ovary, failure of the niche leads to premature differentiation and loss of germline stem cells [2–4]. Conversely, stem cells can become hyperproliferative or tumorigenic if their niche becomes hyperactive, which has been demonstrated in both the *Drosophila* germ cell niche and mammalian hematopoietic stem cell niche [2, 4–6]. Despite progress in understanding the diverse niches of adult stem cells, it has been unclear whether highly potent stem cells—those that are toti- or pluripotent—could be regulated by a similar niche, largely due to the transient nature of these cells during early embryogenesis in most model species.

Unlike many other animals, the planarian *Schmidtea mediterranea* maintains a large pool of pluripotent stem cells throughout adulthood, which the freshwater flatworms use to power whole-body regeneration [7, 8]. Planarian stem cells retain pluripotency throughout life, providing a unique opportunity for study of pluripotent stem cell regulation *in vivo*. Early hints on the stem cell microenvironment came from electron microscopy studies that showed planarian stem cells were found throughout the “parenchyma”—a loosely organized cellular space between organs. Stem cells were often neighbored by differentiated cells called “fixed parenchymal” cells which physically interacted with stem cells through cellular processes and connected to them via gap junctions [9, 10]. More recently, spatial transcriptomic approaches such as MERFISH and SlideSeq revealed that stem cells were found in a diverse cellular microenvironment [11, 12]. These techniques had limited spatial resolution, however, missing narrow cellular processes often seen by fluorescent *in situ* hybridization (FISH) and electron microscopy approaches.

It has also been difficult to pin down local cell types that functionally regulate stem cells. Perturbation of the intestine, which was previously shown be nearby stem cells [12], caused stem cell phenotypes, but it was not clear whether these phenotypes represent local or systemic impacts of the gut [13]. Similarly, planarian muscle plays roles in both axis organization and maintaining the location of stem cells [14, 15], but it is not clear whether muscle-derived factors regulate stem cells locally. On the other hand, secretory cells called hecatonoblasts were identified as a frequent stem cell neighbor through Slide-Seq, but hecatonoblast ablation did not impact stem cell function [12]. One clue emerged from our previous work that revealed *LDLRR3* (*low density lipoprotein receptor related 3*) was often expressed in cells that neighbored stem cells [16]. We showed that *LDLRR3* is also important for optimal brain regeneration, raising the tantalizing possibility that mysterious *LDLRR3^+^* cells could locally influence stem cell function [16].

Given that stem cells are regulated by neighboring cells in other organisms, we first sought to comprehensively and definitively identify differentiated cell types within the parenchymal space and then to test whether stem cell neighbors influence stem cell behavior. We here report unprecedented characterization of an endogenous pluripotent stem cell microenvironment. Interestingly, we observed abundant stem cell – stem cell interactions, suggesting that planarian stem cells could modulate their own microenvironment. We further defined several differentiated cell categories that frequently neighbor stem cells. The most common differentiated neighbor of planarian stem cells is a phagocytic cell type expressing *CTSL2* that we named abraçada cells after the Catalan word for “hugging.” We also identified regional differences in the stem cell microenvironment and investigated how the microenvironment changes after injury and during stem cell differentiation. Based on prior observations that “fixed parenchymal cells” form gap junctions with stem cells, we identified two innexin-encoding genes, *inx8* and *inx9*, expressed in abraçada cells. We find that *inx8* and *inx9* are required for regeneration and animal survival, through modulation of stem cell maintenance and differentiation. We conclude that abraçada cells regulate stem cells non-cell autonomously, with a key mechanism being gap junctions that form between abraçada cells or between abraçada cells and stem cells. Importantly, our findings demonstrate that abraçada cells fulfill both spatial and functional criteria of an endogenous niche for pluripotent stem cells.

## Results and Discussion

### Defining the cellular microenvironment of the planarian pluripotent stem cell

Due to enduring questions about the stem cell microenvironment in planarians (Fig 1A), we first sought to identify differentiated cell types neighboring stem cells. We used single-cell sequencing resources to identify markers of each cell type and pooled these markers together to label the main molecularly-defined categories of differentiated cells in the asexual planarian body: phagocytic/*cathepsin^+^* cells; secretory cells; intestinal cells; neurons; muscle cells; epidermal cells; and protonephridial cells [17, 18](Supp Table 1). We note that we do not use the term “parenchymal” to describe cell identities in this work due to the term’s use to describe separate cell types in the literature. Instead, we reserve the term “parenchymal” to refer to the inter-organ space, which includes planarian stem cells and multiple distinct differentiated cell types. Using double FISH (dFISH), we visualized each cell category along with the stem cell marker *smedwi-1* [19]. For each pool, we determined the percentage of stem cells with one or more labeled neighbors, quantifying cells across 5 body regions and observing cells in all three dimensions (Fig 1B & B’). We counted a cell as a stem cell “neighbor” only when there were no visible unlabeled pixels between the two labeled cells (examples in Fig 1C). This approach was key, because it allowed us to quantify cellular contacts even when stem cell neighbors had complex cell shapes and/or extensive processes, which might have been undercounted or missed by some automated cell counting protocols.

**Figure 1.**
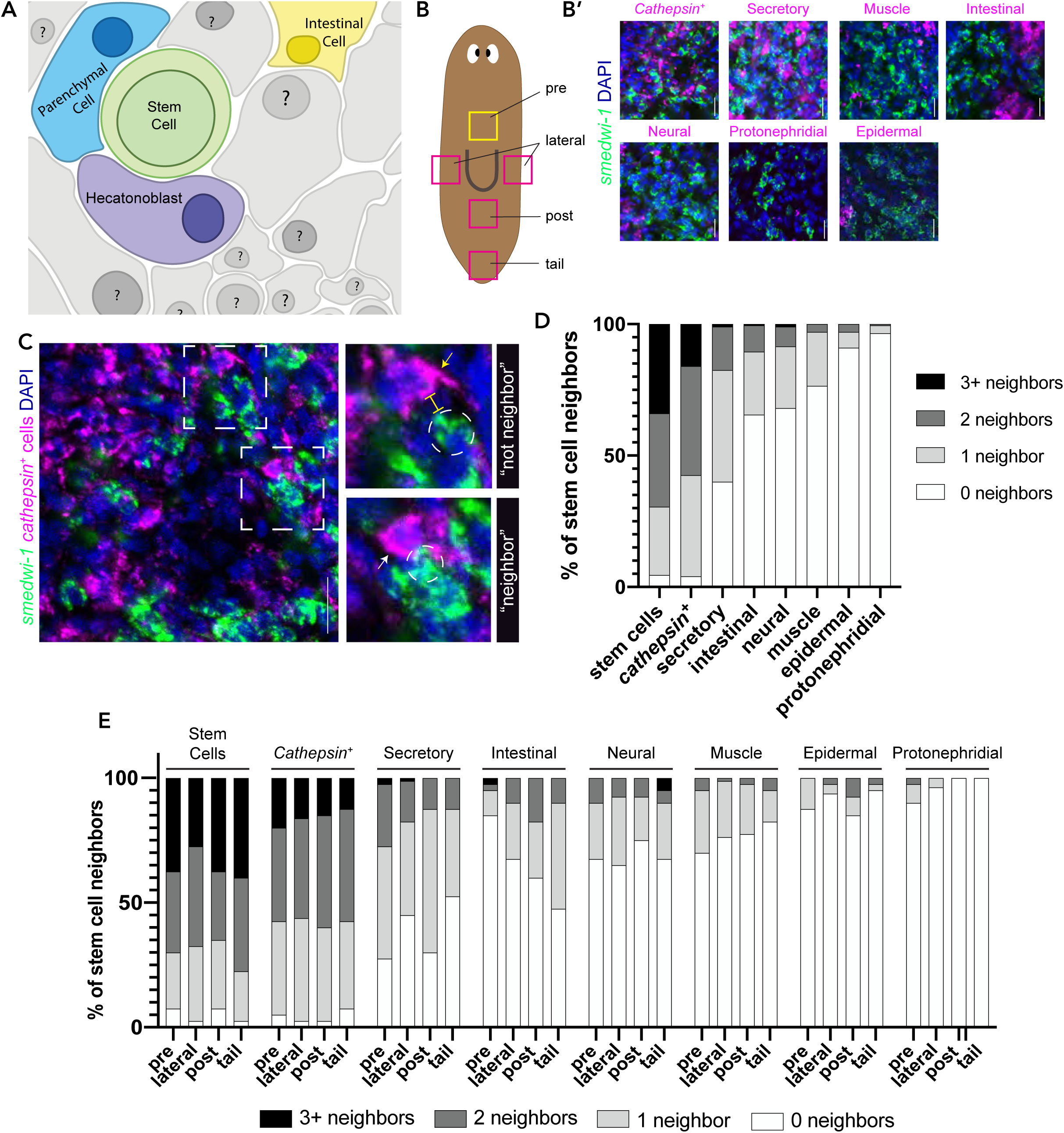
Stem cells neighbor other stem cells and diverse differentiated cell types. (A) Illustration representing the current state of knowledge for the differentiated cell types that are most often found near pluripotent stem cells. (B) Cartoon depicting regions of the planarian imaged in this study. (B’) Representative images of dFISH samples from the pre-pharyngeal region of the planarian, denoted by the yellow box in Fig 1B. In B’, the broad cell category stained is shown in magenta and green staining indicates stem cells marked by *smedwi-1*. Cell nuclei are marked by DAPI, blue. All dFISH images are maximum intensity projections of three 2.5 µm slices. N = 4 for each cell category. (C) An example of stem cell neighbors in a single 2.5 µm slice. The top dashed box and inset illustrates an example of cells that are not “neighbors”: a stem cell (circled) with a *cathepsin^+^* cell nearby (yellow arrow), with a yellow bracket indicating space between the two cells. The bottom dashed box and inset provides an example of a stem cell (circled) with a *cathepsin^+^* neighbor (white arrow). (D-E) Graphs depict the percentage of stem cells that have one or more neighbors for each broad cell category across all regions imaged (D) and with each region shown separately (E). Scale bars: 20 µm.

While we focused on differentiated cell neighbors, the most common neighbor of stem cells was other stem cells (Fig 1D). This observation could indicate that stem cells and their progeny are critical contributors to their own microenvironment. For differentiated cell categories, we found that phagocytic, *cathepsin^+^* cells [17, 18, 20] neighbor stem cells most often, followed by secretory cells and then intestinal cells (Fig 1D). Our data confirm previous studies that showed secretory cells, called hecatonoblasts, and intestinal cell types as frequent stem cell neighbors [11, 12], but our work shows *cathepsin^+^* cells as a key microenvironmental contributor; prior spatial-sequencing studies may have missed this cell type due to complex cell shapes.

Many secretory cell types are enriched around the pharynx, with more sparse numbers in the head and tail. This led us to ask whether the stem cell microenvironment differs regionally in the planarian body. Though stem cells neighbored secretory cells ∼70% of the time immediately anterior and posterior to the pharynx, the frequency of secretory neighbors dropped to ∼50% lateral to the pharynx and in the tail region (Fig 1E). We also observed regional variation for the intestinal cell category, finding that stem cells were more likely to have intestinal neighbors in the tail (Fig 1E). We therefore conclude that the stem cell local environment varies depending on stem cell position in the body, but that *cathepsin^+^* cells remain consistent cellular neighbors throughout the body.

### Stem cell microenvironments during regeneration & differentiation

After injury, planarian stem cells undergo changes in gene expression, cell division, migration, and differentiation [7, 21–27]. We next tested whether the stem cell microenvironment changes after injury. Using the same techniques as above, we examined whether proximity of stem cells to their most frequent neighbors—*cathepsin^+^* and secretory cells (Fig 1D)—changed after injury. We amputated planarians pre- or post-pharyngeally and allowed tissue to regenerate for timepoints up to 3 days, ensuring coverate of the post-injury mitotic bursts at 6 and 48 hours post amputation (hpa) (Wenemoser & Reddien, 2010). During tail or head regeneration, we observed that stem cells neighbored *cathepsin^+^* cells ∼70% of the time at the site of injury a frequency that remained steady or increased slightly as regeneration proceeded (Fig 2B, Supp Fig 1). In contrast, secretory cells neighbored stem cells 45-70% of the time at the site of injury and this frequency decreased significantly as planarians regenerated (Fig 2B, Supp Fig 1). We note that *cathepsin^+^* and secretory cell categories neighbor stem cells at rates slightly different from our maintenance experiment (Fig 1D) likely due to the narrow spatial focus of this experiment.

**Figure 2.**
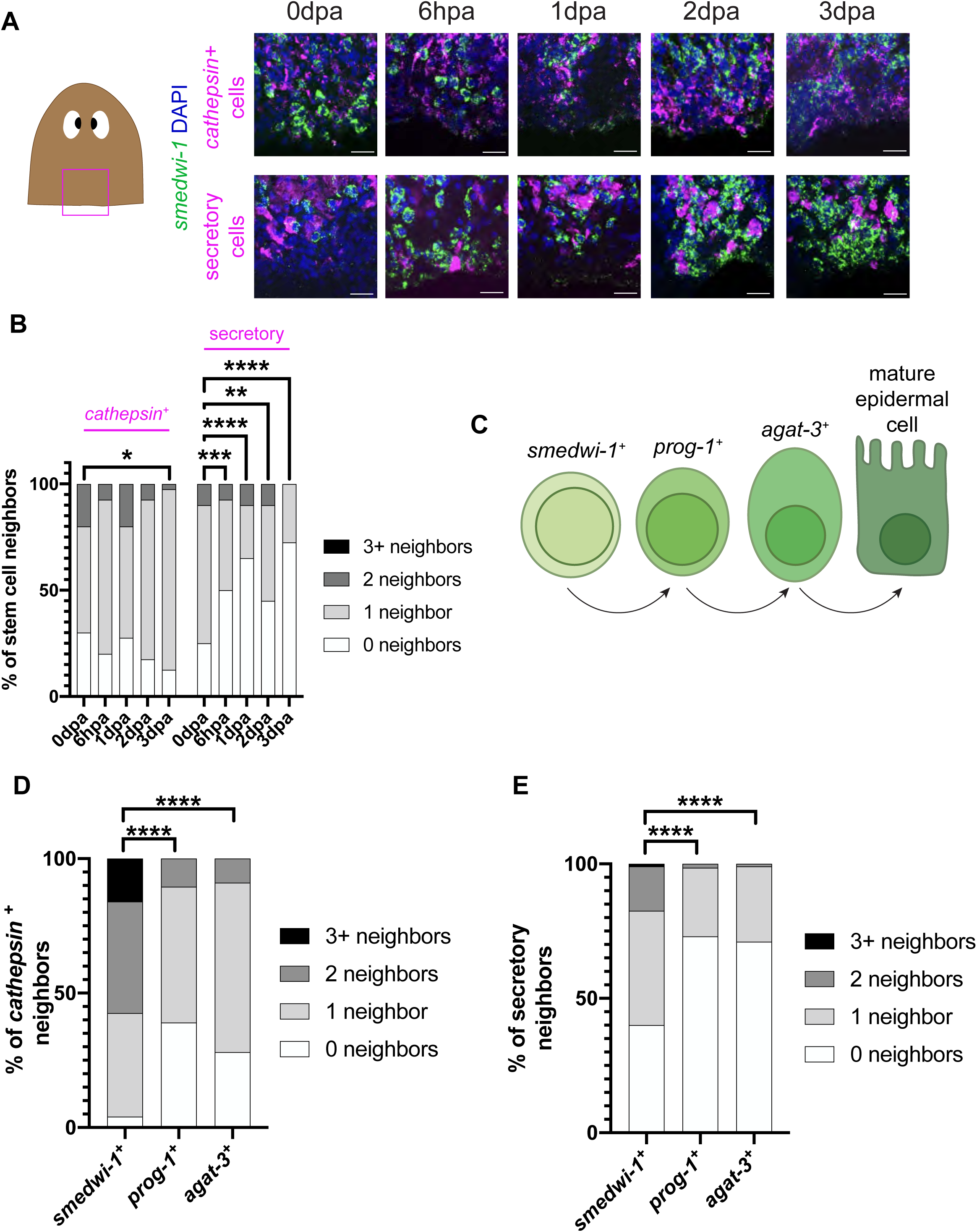
The microenvironment of stem cells changes after injury and upon cell specialization. (A) We examined the stem cell neighbors for *cathepsin^+^* and secretory categories as in Fig 1, but after injury. Images show head fragments regrowing tails over the course of 3 days post-amputation. Images show representative dFISH samples from the injury site, denoted by the magenta box in the cartoon image at left. *Cathepsin^+^* cells (top row) and secretory cells (bottom row) are labeled in magenta, stem cells (*smedwi-1*) in green, and cell nuclei in blue (DAPI). dFISH images are maximum intensity projections of three 2.5 µm slices. N = 4 for each timepoint. Scale bars: 20 µm. (B) Quantification of stem cells with the indicated number of *cathepsin^+^* or secretory neighbors during regeneration. (C) Illustration of markers across the planarian epidermal lineage from pluripotent stem cell to mature epidermal cell [29]. (D & E) Quantification of *smedwi-1^+^* stem cells or differentiating epidermal progenitors (*prog-1^+^* or *agat-3^+^*) with the indicated number of *cathepsin^+^* neighbors (D) or secretory neighbors (E), N = 4 each. Progenitors have significantly fewer of both differentiated neighbor categories. For B and D, significance was determined by comparing cells with at least one neighbor using a Chi-squared test. * p < 0.05; ** p < 0.01; *** p < 0.001; and **** p < 0.0001.

We also wondered how the cellular environment of stem cells changes upon their exit from the pluripotent state and differentiation. In other organisms, distance from the niche or transition through secondary microenvironments promotes differentiation. For example, fly germline stem cells differentiate as they depart from their niche, because they no longer receive cues required for maintenance of the stem cell state [2–4, 28]. For our study, we focused on the well-characterized epidermal lineage, with early (*prog-1*) and late (*agat-3*) markers of specialization [29]. We observed that fate-limited epidermal progenitors marked by *prog-1* and *agat-3* are neighbored by *cathepsin^+^* and secretory cell categories less frequently than *smedwi-1^+^* cells as a group (Fig 2D & 2E). Because we observed a change in the cellular microenvironment correlated with restriction of cell fate, we suspect that either 1) stem cells actively lose contact with their niche upon exiting the pluripotent state *or* 2) stem cell progeny are more prone to differentiation when they stochastically transit away from their optimal niche.

Taken together, our methodology identified *cathepsin^+^* cells as stem cells’ most proximal differentiated neighbor across the entire body and after injury, but we note that our finding does not exclude the possibility that stem cells are impacted by other local cell types and long-range signals supplied by cells outside of the niche. Within the planarian field, there is already evidence that cell categories like muscle and neurons impact stem cell function despite the relative distance that we demonstrated in our work. For example, muscle supplies positional cues for the AP axis through position control genes that influence stem cell fate choices [14, 15, 30, 31]. Regardless, *cathepsin^+^* cells neighbor stem cells at a high rate in both maintenance and regeneration, so we proceeded to investigate the *cathepsin^+^* cell category as a potential component of to the local pluripotent stem cell niche.

### Narrowing in on key cellular neighbors

The phagocytic/*cathepsin^+^* category comprises multiple distinct cell types and/or states, including glial cells, pigment cells, and undefined cell types, united by transcriptomic similarities in single cell sequencing studies (Fig 3A) [17, 18]. We identified markers that label narrower subsets of *cathepsin^+^* cell types and states (Fig 3A). We did not examine glial cells or pharyngeal *cathepsin^+^* cells (*TTPA^+^*) due to exclusion of stem cells from the neuropil and the pharynx respectively [17, 18] (Fig 3A). We then performed additional dFISH experiments to determine which *cathepsin^+^* subtypes are most frequently found in the stem cells’ local environment. Our data showed that stem cells are neighbored by cells expressing *cathepsin L2 (CTSL2)* almost as frequently as they are neighbored by cells of the entire phagocytic cell category (Fig 3B-C). *LDLRR3,* which we had previously characterized, also labels a group of several cell types that neighbor stem cells 70% of the time (Fig 3B, D). We noted broad but incomplete coexpression of *LDLRR3* and *CTSL2* (Supp Fig 2). *Other cathepsin^+^/*phagocytic cell types neighbor stem cells at varying rates, including pigment cells (*HMBS^+^*) and uncharacterized cell types marked by *PTPRG*, *PTPRT*, or *dd5960* (Fig 3B, E-I).

**Figure 3.**
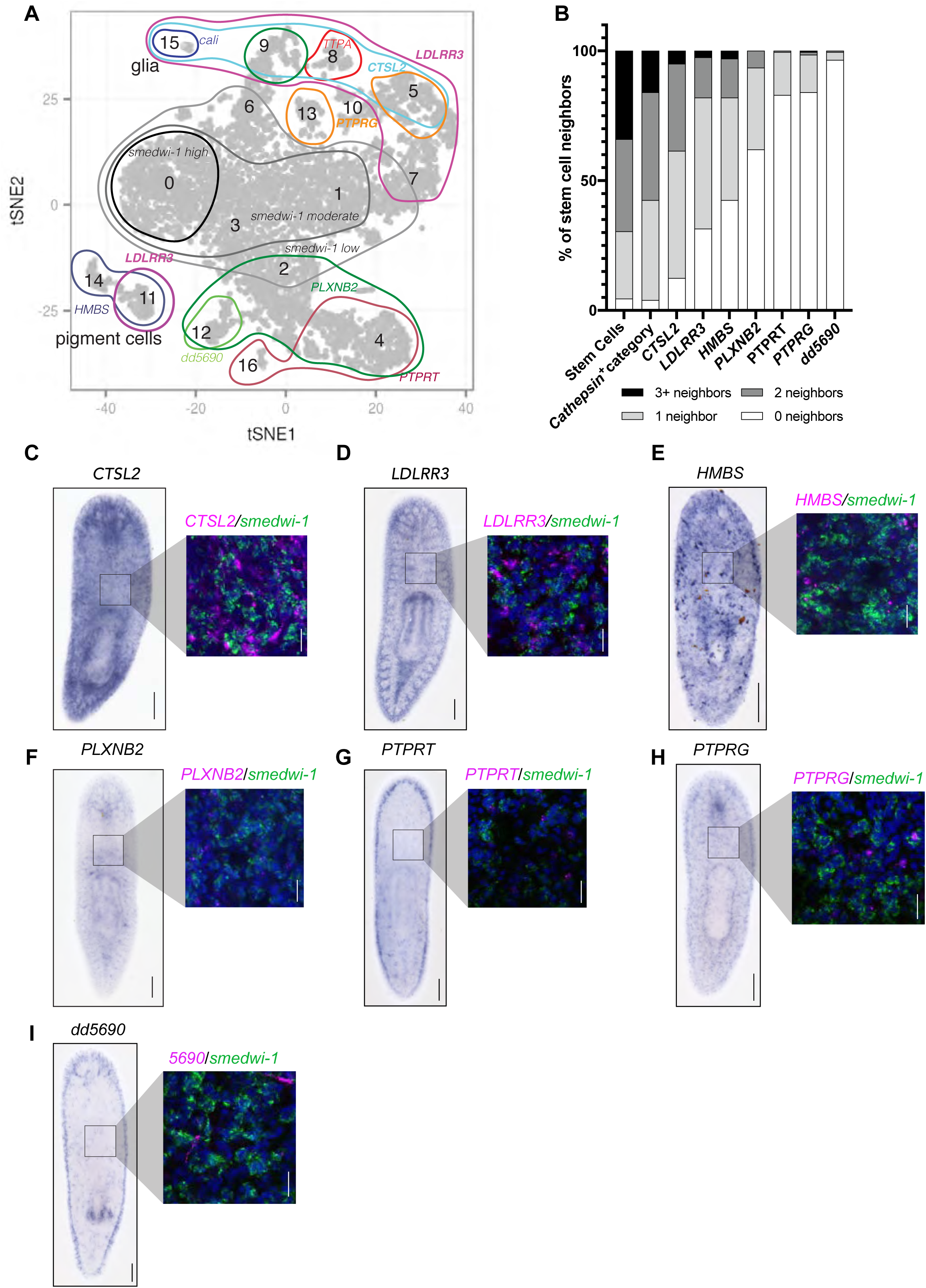
Stem cells neighbor *CTSL2^+^* abraçada cells. (A) *Cathepsin^+^* cluster from [17] showing expression of more restricted markers. (B) Quantification of stem cells with cells expressing each marker as neighbors. (C-I) *In situ* hybridization (ISH) for each *cathepsin^+^* cell type with representative inset of dFISH samples from the pre-pharyngeal region of the planarian, denoted by the black outlined box. Magenta is used for each marker; green labels stem cells (*smedwi-1*), and DAPI is in blue. All dFISH images are maximum intensity projections of three 2.5 µm slices, N = 4. ISH scale bars: 200 µm; dFISH scale bars: 20 µm.

Interestingly, *CTSL2* marks four distinct *cathepsin^+^* cell types or states in sequencing atlases (*cathepsin^+^* “sub-clusters” 5, 8, 9 and 15 in the atlas created by Fincher, et al) [17]. Two *CTSL2*^+^ subclusters [pharyngeal *TTPA^+^* cells (Fincher sub-cluster 8) and glia (Fincher sub-cluster 15)] are not present in the parenchymal space in which stem cells reside [17]. Because most *cathepsin^+^* cell types are currently uncharacterized, we named parenchymal *CTSL2^+^* cells “abraçada cells” after the Catalan translation of “hugging,” based on our observation that parenchymal *CTSL2^+^* cells are often immediately adjacent to stem cells, occasionally enwrapping or “hugging” them with cellular processes. We were unable to identify markers to refine our molecular definition of abraçada cells beyond this point due to the transcriptional similarity of remaining sub-clusters (Fincher *cathepsin^+^* 5 and 9). We conclude that remaining heterogeneity within abraçada cells could indicate variable cell states and/or identities of the stem cell niche under different conditions or in different parts of the planarian body.

### Abraçada cells function as a stem cell niche

One essential feature of a bona fide stem cell niche is that it impacts stem cell function. Previous research showed that planarian stem cells are in physical contact with neighboring differentiated cell types through gap junctions [9]. In invertebrates, gap junctions are composed of proteins called innexins [32] and some innexins play documented roles in planarian polarity and stem cell function [33, 34]. We reasoned that abraçada cells could function as a niche-providing cell by using gap junctions to facilitate stem cell function. Using a transcriptomic atlas [17], we identified two innexin-encoding genes homologous to *Dugesia japonica inx8* and *inx9* [34] (Supp Table 2-3), expressed in *cathepsin*^+^ cell types in a pattern very similar to *CTSL2* and *LDLRR3* and not expressed in stem cells themselves (Supp Fig 3A-C). To test whether innexins, and thus abraçada cells, promote proper stem cell function, we knocked down *inx8* and *inx9* with RNA interference (RNAi). After RNAi, we amputated animals pre-pharyngeally and allowed animals to regenerate for 7 days. After *inx8(RNAi)* or *inx9(RNAi),* we observed a statistically significant reduction in blastema size relative to body size (Fig 4A-B), showing that broad regenerative capacity is altered. We found an even more striking regenerative failure upon double RNAi of *inx8* and *inx9*, quantified via reduction in regenerated brain size (Fig 4C-D).

**Figure 4.**
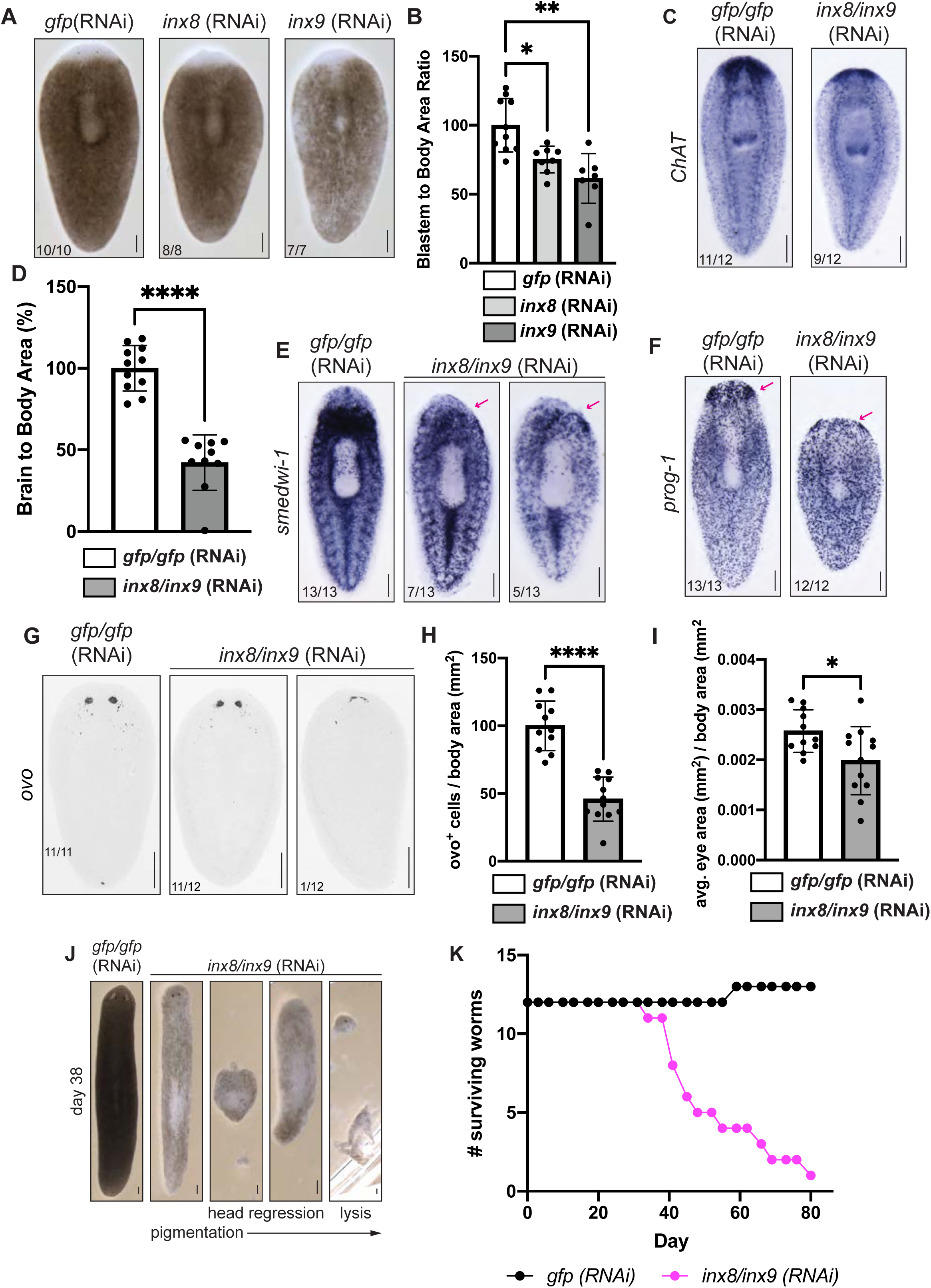
Innexins mediate a non-cell autonomous effect of abraçada cells on stem cells. (A) RNAi experimental worms regenerating heads. Planarians are shown here with unpigmented blastemas visible. (B) Quantification of blastema-to-body ratio of animals from A normalized so that gfp(RNAi) results are set to 100%. One-way ANOVA with Dunnett’s multiple comparison ; * p = 0.0354; ** p = 0.0013. (C) ISH of *gfp;gfp*(RNAi) and *inx8(RNAi);inx9(RNAi)* animals after head regeneration stained for *choline acetyltransferase* (*ChAT*). (D) Quantification of brain-to-body area ratios in C, normalized to percentage of controls. Unpaired t-test: **** p < 0.0001. (E-F) *In situ* hybridization of *gfp;gfp*(RNAi) and *inx8(RNAi);inx9(RNAi)* worms after head regeneration stained for *smedwi-1* and *prog-1*. Magenta arrows indicate regions of reduced expression. (G) *gfp;gfp*(RNAi) and *inx8(RNAi);inx9(RNAi)* animals after head regeneration with FISH for *ovo* and presented as an inverse-color image. (H-I) Quantification of the number of *ovo^+^* eye progenitors normalized to body area (H) & eye size normalized to body area (I). Unpaired t-tests were performed: **** p < 0.0001; * p = 0.0222. (J) Long term phenotypes for *gfp;gfp*(RNAi) and *inx8(RNAi);inx9(RNAi)* animals. We saw variation of phenotypes in the innexin double knockdowns, starting on the left with reduced pigmentation, head regression & lysis phenotypes. (K) Survival curve of *gfp;gfp*(RNAi) and *inx8(RNAi);inx9(RNAi)* animals. Scale bars: 200 µm.

We next asked whether perturbation of innexin genes impacts regeneration by impacting stem cell behavior. First, we tested whether innexin perturbation leads to a decline in the stem cell population itself. Indeed, we observed a visible reduction in *smedwi-1^+^* cells throughout the body that ranged from modest to striking (Fig 4E). We also noted less *smedwi-1* enrichment near the site of injury compared to controls (Fig 4E). We reasoned that a failure of local enrichment might be due to altered changes in proliferation, but we did not see any change in mitotic cells between control and experimental worms (Supp Fig 4). We next examined whether stem cell progeny were depleted, using the epidermal and eye lineages, marked by *prog-1* and *ovo* respectively [29, 35, 36]. While *prog-1^+^* cells appeared unaffected in maintained tissue, we saw a decrease in *prog-1^+^* enrichment within the regenerating tissue of experimental worms (Fig 4F). After perturbation of *inx8* and *inx9*, we also noted a decrease in the number of eye progenitors marked by *ovo* (Fig 4G-H) that ultimately led to failure to regenerate eyespots of normal size (Fig 4G, I). Taken together, our results support a model in which innexin perturbation impacts the ability of stem cells to respond robustly to injury to produce adequate progeny for a pro-regenerative outcome.

Given the impact of innexins on stem cell function and the importance of stem cells for planarian survival, we also asked whether planarian innexins expressed in abraçada cells were essential to animal survival. After 13 days, we observed pigmentation phenotypes in *inx8(RNAi);inx9(RNAi)* animals. After 38 days, we saw lysis and severe head regression (Fig 4J). Animals began to die after 34 days of RNAi and all worms, except one, died over the course of an 80-day RNAi treatment (Fig 4K). The typical survival time for planarians after lethal irradiation is around 2-3 weeks, which is shorter than we observe in our survival paradigm. The timing of animal death and our prior data are not consistent with abraçada innexins being absolutely and immediately required for stem cell viability. Instead, our data are more consistent with stem cells failing to adequately self-renew and replace cells lost to homeostatic turnover, resulting in eventual head regression and lysis.

Taken together, our data support a model in which functional gap junctions in abraçada cells non-cell autonomously promote nearby stem cell differentiation, mobilization after injury, and maintenance. We further note that *LDLRR3* was originally identified as a wound-induced gene that itself impacts regeneration [16, 22]. *LDLRR3* encodes a putative secreted protein that may provide a second, injury-enhanced mechanism through which abraçada cells influence stem cell function. We have therefore demonstrated in two distinct ways that perturbation of genes expressed in abraçada cells leads to deleterious effects on stem cell function and regeneration.

We conclude that abraçada cells serve as an endogenous niche for pluripotent stem cells. In the future, we plan to identify a strategy with which to eliminate abraçada cells, which will allow us to capture the full range of functions for this novel cell type. Our investigation of the nature and function of an *in vivo* pluripotent stem cell niche provides fundamental information that can improve culture of planarian stem cells *in vitro*, which is currently not possible [37] and may eventually improve mammalian pluripotent stem cell induction and culture. Based on our work we also propose that innexins may also prove to be an attractive drug target for anti-helminthic therapies, especially if stem cells in parasitic flatworms respond to a similar niche. More broadly, our endogenous portrait of the pluripotent stem cell microenvironment could lead to discovery of new molecules and pathways that regulate pluripotency.

## Limitations of the study

We note that mRNA is not always localized ubiquitously throughout a cell, so it is possible that our quantification of cellular neighbors through FISH is overly conservative; future work to identify membrane markers of abraçada cell types and to develop immuno-electron microscopy will allow us to more accurately identify and define cellular contacts within the planarian niche. We also conclude that some heterogeneity of cell type and state may still exist within our definition of abraçada cells. Future work will be needed to determine the specific nature of heterogeneity within the cell category, including whether abraçada cells change state after injury.

## Acknowledgements

We thank all current members and alumni of the Roberts-Galbraith lab for technical training and idea development. Specifically, we thank Kendall Clay, Taylor Medlock-Lanier, Macey Wilson, Christina Endara-Arnold, Jazmine Dent, and Camila Gomes Da Silva for providing comments on this manuscript. We thank the University of Georgia Biomedical Microscopy Core and Dr. Muthugapatti Kandasamy for training and help with fluorescent imaging. We thank Dr. Shannon Quinn for advice on ongoing efforts to quantify cell neighborhoods. We acknowledge and thank the sources of funding that made this work possible: National Science Foundation CAREER award to Dr. Roberts-Galbraith (IOS-1942822), National Institute of Health Genetics Training Grant support of Skylar Settles (T32GM142623) and University of Georgia Center for Undergraduate Research Opportunities support of Kathleen Miller. RRG is also supported by the Lois K. Miller Professorship of Cellular Biology at UGA. Finally, we thank White Oak Pastures (Bluffton, GA) for providing beef liver for our planarians.

## Materials and Methods

### Animal Husbandry

For all experiments, the asexual strain of *Schmidtea mediterranea* was used (CIW4) [38]. Planarians were maintained in a large recirculation enclosure where they were kept in 1x Montjuïc salts [38]. Before experiments, planarians were housed in 9 cup Ziploc containers filled with approximately 1 liter of 1x Montjuïc salts and kept at 18°C in the dark. Biweekly, planarians were fed pureed beef liver obtained from White Oak Pastures, GA, USA and were treated with gentamicin sulfate (50 µg/mL final concentration) as needed which was approximately once per month. Worms were starved 1-2 weeks before experiments and were starved for 7 days prior to amputations. Amputations were performed with a scalpel either anterior (pre-pharyngeally) or posterior (post-pharyngeally) to the pharynx.

### Identification of cell type markers

A published whole-body planarian transcriptomic atlas was used to identify markers of each cell type that were not expressed in stem cells or other cell categories [17] (Supp Table 1). When examining differentiated cell types from single-cell sequencing data, we used the absence of *smedwi-1^+^* mRNA as an indicator of a differentiated cell type. Supp Table 1 shows each gene that was used as a marker and indicates the cell types (“subclusters”) and cell categories (“main cluster”) they label, using [17] as a reference. Differentiated pharyngeal cells were not included in our study, because no stem cells are present within this tissue. For a small number of cell clusters or types, we were unable to identify a unique transcript with which to complete labeling. We hypothesize that the cell types without unique transcripts may represent transitional states.

### Identification of genes and cloning

Sequences for genes of interest were determined using PlanMine3.0 [39]. All primers used to amplify genes are available in Supp Table 1 and 4 and were designed using Primer3Plus [40] to amplify a 400-1000 bp sequence. Gene fragments were cloned from cDNA following methods outlined in [41]. Briefly, asexual *S. mediterranea* cDNA was synthesized with an iSCRIPT kit (BioRad) and primers were used to amplify the gene fragments. Amplified fragments were ligated into a pJC53.2 vector that had been digested with EAM1105I (New England Biolabs). Ligation products were transformed into *E. coli* with kanamycin selection and isolated with Zyppy Plasmid Miniprep Kit (Zymo). All plasmids were sequenced to ensure the gene was cloned properly (Azenta) and to determine the orientation of gene insertion. Genes of interest were amplified from pJC53.2 vectors using a T7 primer (Supp Table 4) and cleaned using a DNA Clean and Concentrator Kit (Zymo) for downstream methods.

### *In situ* hybridization

Asexual planarian colorimetric and fluorescent *in situ* hybridization were performed as outlined in [42], protocols adapted from [43]. Briefly, single-stranded antisense riboprobes were transcribed with Digoxigenin-11-UTP (Roche) or Fluorescein-12-UTP (Roche) following steps outlined in [41]. For colorimetric *in situ* hybridization, riboprobes were synthesized using Digoxigenin-11-UTP (Roche) and were detected using anti-digoxigenin Fab fragments conjugated to alkaline phosphatase (Roche). We visualized antibodies in colorimetric *in situ* hybridization with BCIP (Roche) and NBT (Roche) in an alkaline phosphatase buffer. For fluorescent *in situ* hybridization, riboprobes made using Fluorescein-12-UTP (Roche) were detected using anti-fluorescein Fab fragments conjugated to peroxidase (Roche), while riboprobes created with Digoxigenin-11-UTP (Roche) were detected using anti-digoxigenin Fab fragments conjugated to peroxidase (Roche). We developed fluorescent signal with fluor-tyramide FAM or fluor-tyramide TAMRA for fluorescein-12-UTP and digoxigenin-11-UTP riboprobes respectively and quenched peroxidases with 0.1M sodium azide (ThermoFisher Scientific) between antibodies

### Immunofluorescence

The immunofluorescence protocol used was adapted from published protocols [44, 45] and summarized in [46]. Briefly worms were killed in 2% HCl and phosphorylated histone 3 was observed with the primary antibody rabbit-anti-phospho-histone H3 (Ser10) using a 1:1600 dilution (Cell Signaling Technology). To obtain more specific signal amplification the secondary antibody, goat-anti-rabbit IgG-HRP, was used at a 1:1000 dilution (Invitrogen, ThermoFisher Scientific) and the fluorescent signal was developed with fluor-tyramide FAM.

### Sample imaging

Colorimetric samples were mounted in 80% glycerol and imaged with an Axiocam 506 camera on a Zeiss Axio Zoom V.16 microscope using ZEN 2.3 Pro software. Fluorescent samples were mounted in VectaShield and imaged on a Zeiss 880 LSM confocal microscope with Zen Black 2.3 SPI software. Fluorescent samples from all stem cell neighbor experiments were imaged using Plan-Apochromat 40x/0.8 oil immersion in 2.5 µm interval z-stack slices and the first and last boundaries were determined based on fluorescent signal (Figures 1-3). Fluorescent samples probing for *ovo* were imaged using EC Plan-Neofluar 10x/0.30 M27 (Fig 4G).

### Quantification of stem cell neighbors

For homeostasis experiments, 5 regions of each planarian were imaged to acquire representative images for quantification. The 5 regions imaged include anterior and posterior to the pharynx, to the left and right of the pharynx, as well as toward the end of the tail. For the regeneration timeline, stem cells were imaged at the injury site. Manual quantification of stem cell neighbors was accomplished using the cell counter plugin on Fiji [47]. With only the stem cell channel visible (FAM or green channel in all experiments), 10 stem cells were selected in each region. After cell selection, all channels were merged and the differentiated cells that directly neighbored each stem cell were quantified. We were careful to examine each stem cell through space, evaluating neighbors in z slices above, through, and below each cell. Manual quantification was performed conservatively and if any space was visible between a stem cell and differentiated cell, it was not quantified as a “neighbor”. For each experiment, 4 worms were imaged. Thus, the total number of stem cells quantified was 200 per cell type or cell category.

### RNAi experiments

dsRNA was synthesized from pJC53.2 clones as previously described [48]. dsRNA concentration was quantified from a 1:10 dilution of each dsRNA sample through gel electrophoresis. The band was quantified relative to a known band of 1 kb Hyperladder to determine dsRNA concentration. For RNAi experiments, 10-12 worms were included in a 60 mm Petri dish and fed 5 µg of each dsRNA mixed with 30 µl pureed beef liver and 1 µl green food dye to confirm that the worms ate the liver mixture. For double RNAi experiments, we included 5 µg of each mRNA target for a total of 10 µg of dsRNA. Negative controls were fed dsRNA matching *Aequorea victoria green fluorescent protein (GFP),* which is not found in the planarian genome. Experimental worms were allowed to eat for 2-3 hours. After feeding, the liver mixture was removed and worms were rinsed 4 times and transferred to fresh 60 mm Petri dishes with fresh Montjuïc salts. The regeneration RNAi paradigm followed (for all experiments except Fig 4J-K) was 5 feedings 3 days apart, followed by a 7-day starvation period then amputation. After amputation, experimental worms were allowed to regenerate for 7 days before being fixed for downstream experiments. The survival RNAi paradigm (Fig 4J-K) involved feeding control and experimental worms every 3-4 days for 80 days. Survival was monitored prior to each feeding.

---

**Supplemental Figure 1.**
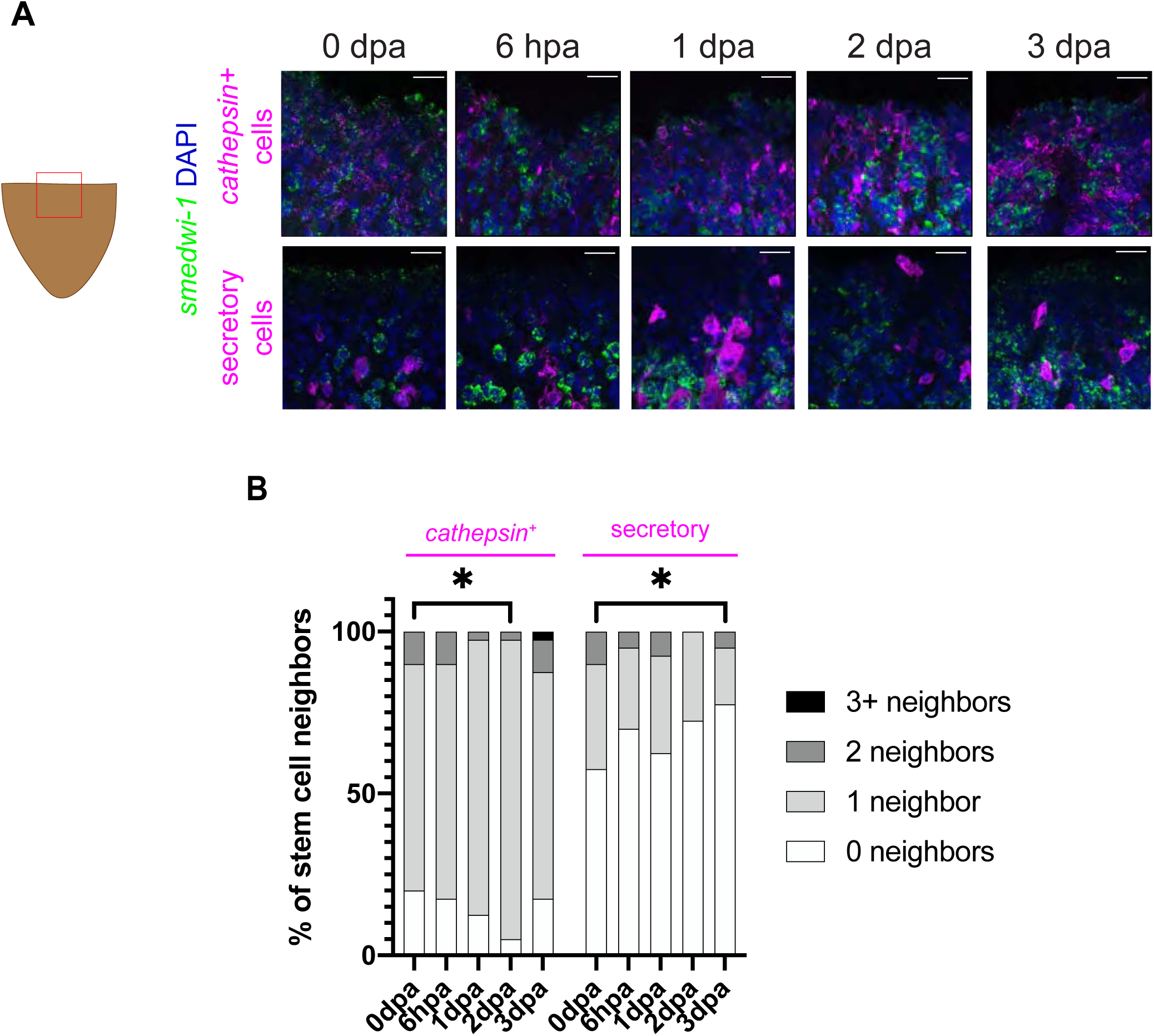
(A) Stem cell neighbors for *cathepsin^+^* and secretory categories after injury. We here show tails regrowing heads over the course of 3 days post-amputation (dpa). Images show representative dFISH imaged at the injury site, denoted by the magenta box in the cartoon image on left. All images are oriented in the same direction as the cartoon image with tail pointing down. *Cathepsin^+^* cells (top row) or secretory cells (bottom row) are labeled in magenta, stem cells marked by *smedwi-1* are labeled in green, and cell nuclei marked by DAPI are in blue. All dFISH images are maximum intensity projections of three 2.5 µm slices. N = 4 for each timepoint. Scale bars: 20 µm. (B) Quantification of the percentage of stem cells with the indicated number *cathepsin^+^* or secretory neighbors throughout regeneration. Significance was determined through Chi-squared tests, comparing cells with at least one neighbor. * p < 0.05; ** p < 0.01; *** p < 0.001; and **** p < 0.0001.

**Supplemental Figure 2.**
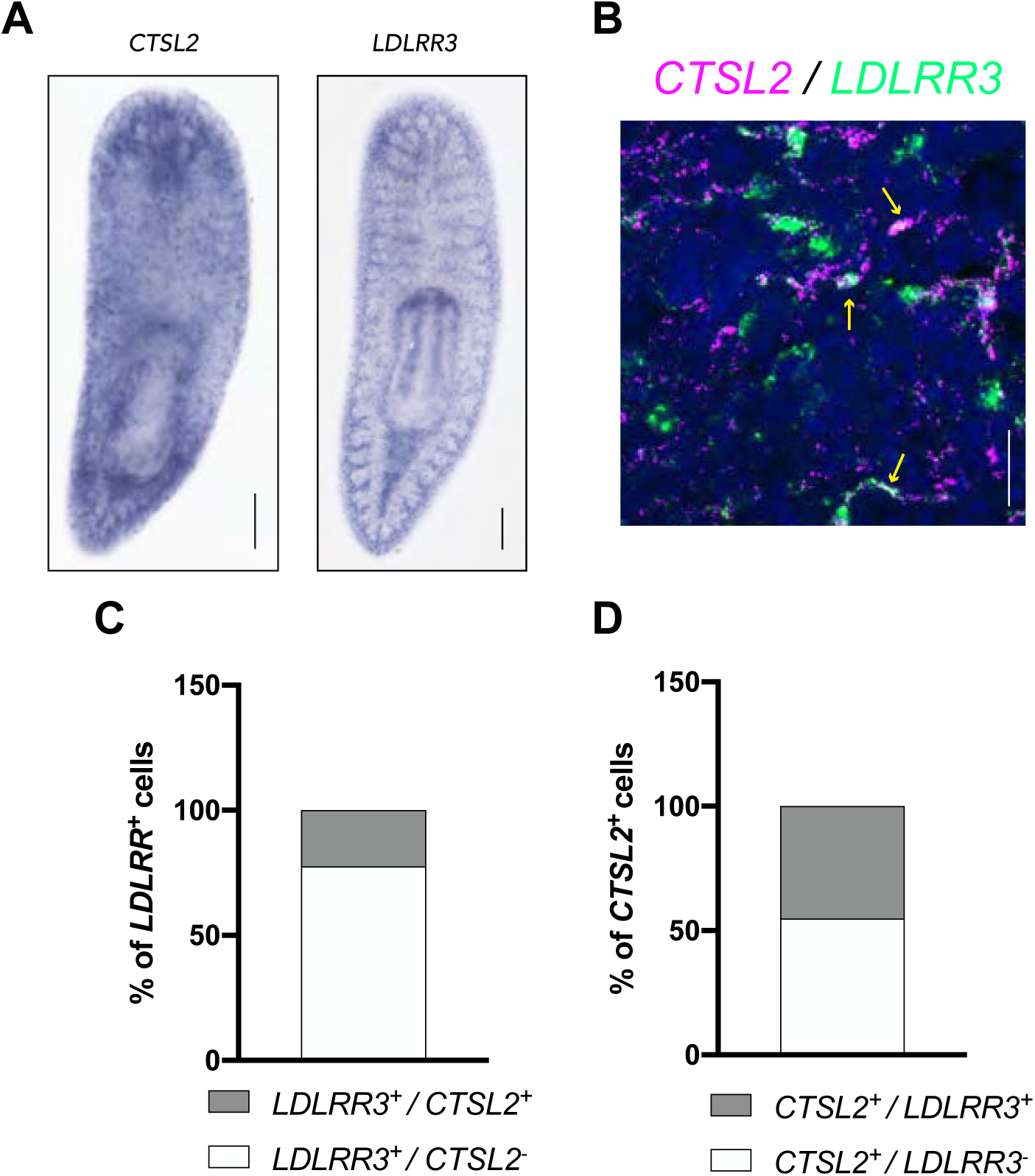
(A) ISH expression of *CTSL2* and *LDLRR3.* Scale bar: 200 µm. (B) Representative dFISH image in the pre-pharynx region showing co-expression. *CTSL2* is shown in magenta and *LDLRR3* is shown in green, with DAPI shown in blue. Yellow arrows indicate cells expressing the two markers. Scale bar: 20 µm and N = 4. (C) Coexpression quantification of the percentage of *LDLRR3* cells that are either *CTSL2^+^* or *CTSL2^-^* (D) Coexpression quantification of the percentage of *CTSL2*^+^ cells that are either *LDLRR3^+^* or *LDLRR3^-^*.

**Supplemental Figure 3.**
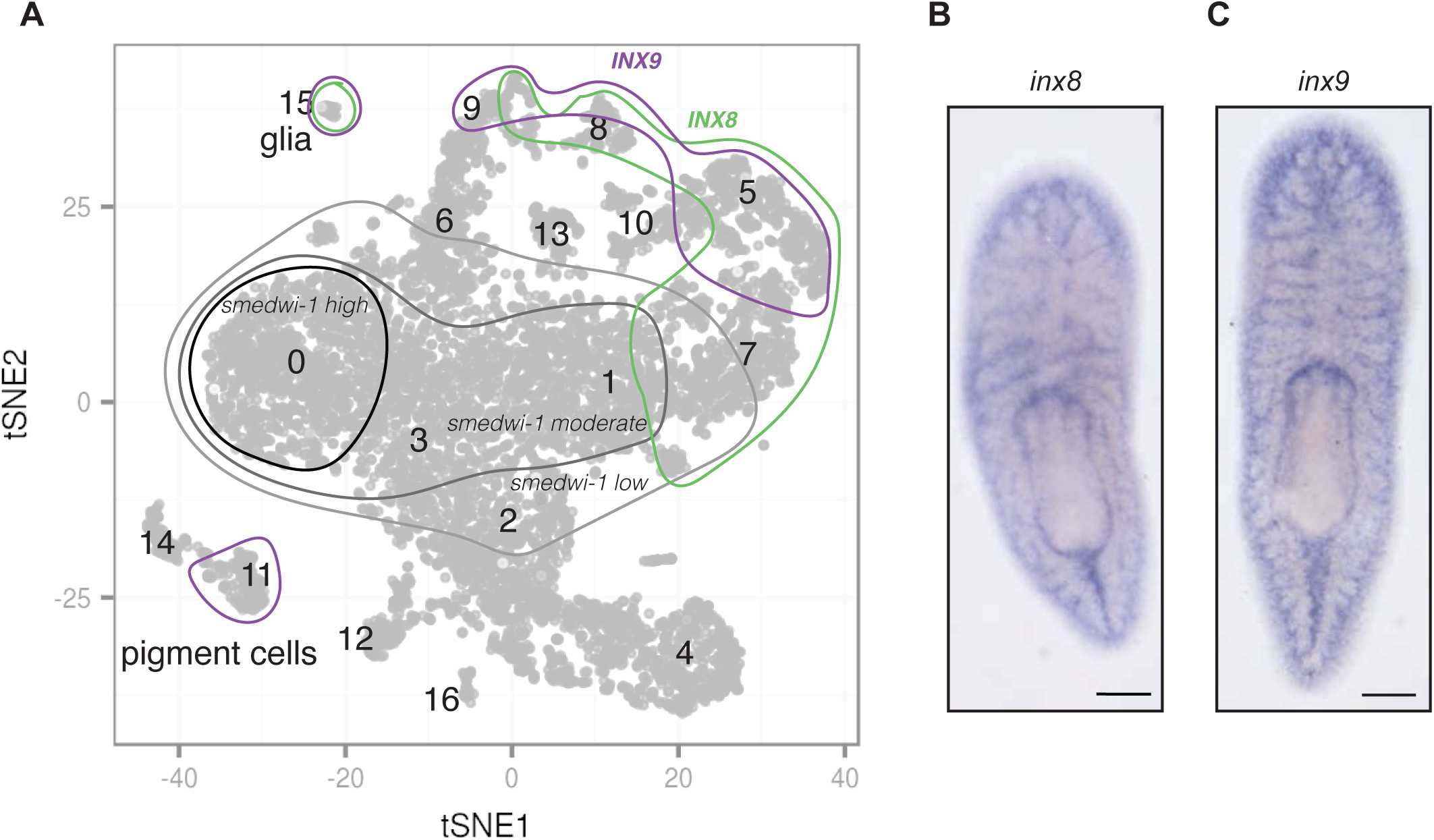
Cathepsin^+^ cluster from [17] showing sub-clusters that express *innexins 8* and *9*. (B & C) ISH expression patterns of *inx8* (B) and *inx9* (C). Scale bar: 200 µm.

**Supplemental Figure 4.**
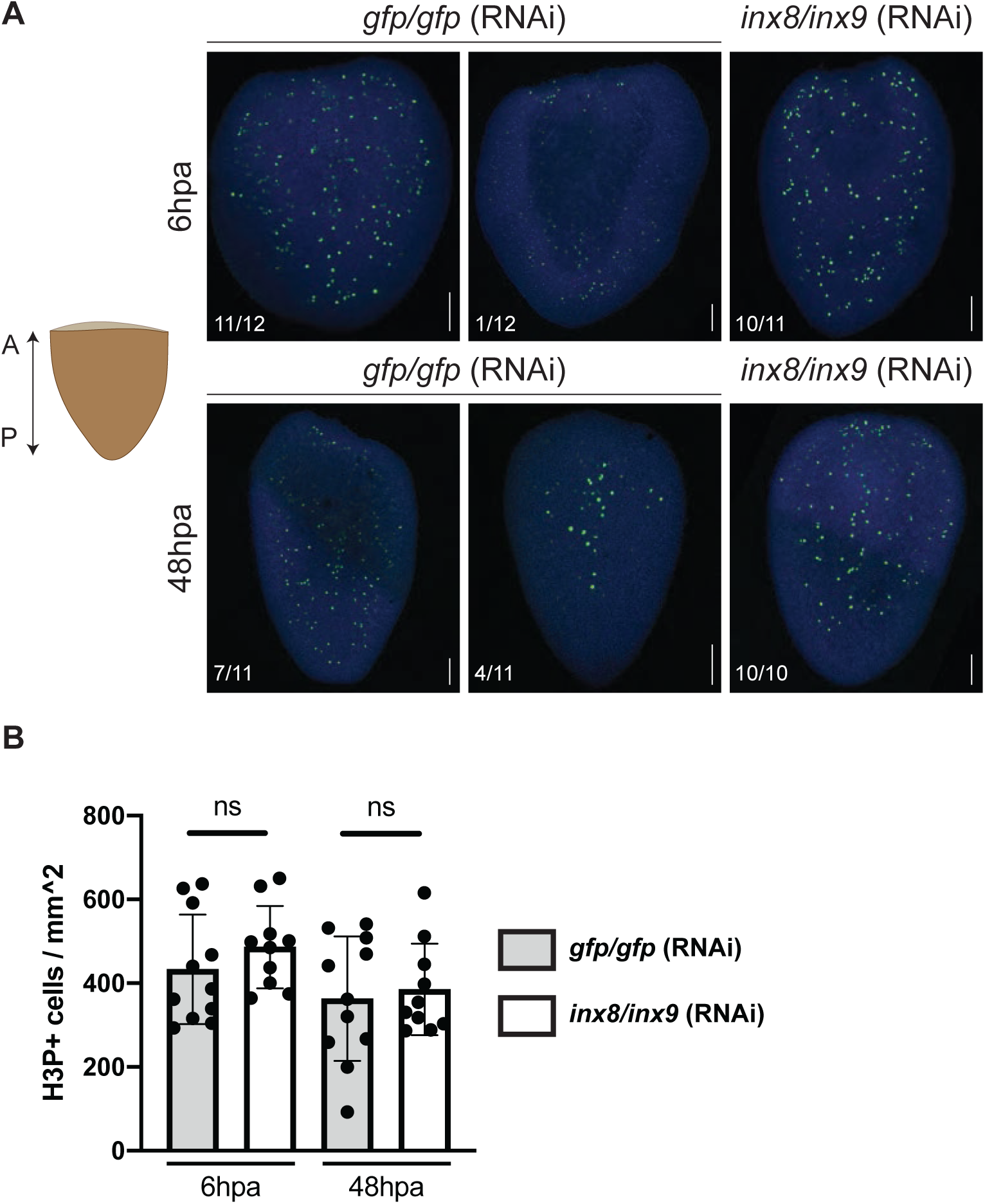
(A) Mitotic activity after innexin RNAi shown by H3P labeled in green, DAPI shown in blue. Images are shown of tails regenerating heads at 6 hpa (top row) or 48 hpa (bottom row). Scale bar: 100µm. (B) Quantification of number of H3P+ cells throughout the entire body. An unpaired t-test was performed to assess statistical significance.

## References

1. Schofield, R., The relationship between the spleen colony-forming cell and the haemopoietic stem cell. Blood cells, 1978. 4(1-2): p. 7–25.

2. Song, X., et al., Bmp signals from niche cells directly repress transcription of a differentiation-promoting gene, bag of marbles, in germline stem cells in the Drosophila ovary. Development, 2004. 131(6): p. 1353–1364.

3. Song, X. and T. Xie, DE-cadherin-mediated cell adhesion is essential for maintaining somatic stem cells in the Drosophila ovary. Proceedings of the National Academy of Sciences, 2002. 99(23): p. 14813–14818.

4. Xie, T. and A.C. Spradling, decapentaplegic Is Essential for the Maintenance and Division of Germline Stem Cells in the Drosophila Ovary. Cell, 1998. 94(2): p. 251–260.

5. Calvi, L.M., et al., Osteoblastic cells regulate the haematopoietic stem cell niche. Nature, 2003. 425(6960): p. 841–6.

6. Zhang, J., et al., Identification of the haematopoietic stem cell niche and control of the niche size. Nature, 2003. 425(6960): p. 836–41.

7. Baguna, J., E. Saló, and C. Auladell, Regeneration and pattern formation in planarians. III. Evidence that neoblasts are totipotent stem cells and the source of blastema cells. Development, 1989. 107: p. 77–86.

8. Wagner, D.E., I.E. Wang, and P.W. Reddien, Clonogenic Neoblasts Are Pluripotent Adult Stem Cells That Underlie Planarian Regeneration. Science, 2011. 332(6031): p. 811–816.

9. Hori, I., Role of fixed parenchyma cells in blastema formation of the planarian Dugesia japonica. The International Journal of Developmental Biology, 1991. 35(2): p. 101–108.

10. Pedersen, K.J., Studies on the nature of planarian connective tissue. Zeitschrift für Zellforschung und Mikroskopische Anatomie, 1961. 53(5): p. 569–608.

11. Park, C., et al., Fate specification is spatially intermingled across planarian stem cells. Nature Communications, 2023. 14(1): p. 7422.

12. Benham-Pyle, B.W., et al., Planarians employ diverse and dynamic stem cell microenvironments to support whole-body regeneration. bioRxiv, 2023: p. 2022.03.20.485025.

13. Forsthoefel, D.J., et al., Cell-type diversity and regionalized gene expression in the planarian intestine. eLife, 2020. 9: p. e52613.

14. Witchley, Jessica N., et al., Muscle Cells Provide Instructions for Planarian Regeneration. Cell Reports, 2013. 4(4): p. 633–641.

15. Cote, L.E., E. Simental, and P.W. Reddien, Muscle functions as a connective tissue and source of extracellular matrix in planarians. Nature Communications, 2019. 10(1): p. 1592.

16. Roberts-Galbraith, R.H., J.L. Brubacher, and P.A. Newmark, A functional genomics screen in planarians reveals regulators of whole-brain regeneration. eLife, 2016. 5: p. e17002.

17. Fincher, C.T., et al., Cell type transcriptome atlas for the planarian *Schmidtea mediterranea*. Science, 2018. 360(6391): p. eaaq1736.

18. Plass, M., et al., Cell type atlas and lineage tree of a whole complex animal by single-cell transcriptomics. Science, 2018. 360(6391): p. eaaq1723.

19. Reddien, P.W., et al., SMEDWI-2 is a PIWI-like protein that regulates planarian stem cells. Science, 2005. 310(5752): p. 1327–30.

20. Scimone, M.L., et al., foxF-1 Controls Specification of Non-body Wall Muscle and Phagocytic Cells in Planarians. Current Biology, 2018. 28(23): p. 3787–3801.e6.

21. Wurtzel, O., et al., A Generic and Cell-Type-Specific Wound Response Precedes Regeneration in Planarians. Developmental Cell, 2015. 35(5): p. 632–645.

22. Wenemoser, D., et al., A molecular wound response program associated with regeneration initiation in planarians. Genes Dev, 2012. 26(9): p. 988–1002.

23. Saló, E. and J. Baguñà, Stimulation of cellular proliferation and differentiation in the intact and regenerating planarian Dugesia(G) tigrina by the neuropeptide substance P. Journal of Experimental Zoology, 1986. 237(1): p. 129–135.

24. Newmark, P.A. and A. Sánchez Alvarado, Bromodeoxyuridine Specifically Labels the Regenerative Stem Cells of Planarians. Developmental Biology, 2000. 220(2): p. 142–153.

25. Ito, H., et al., Epimorphic regeneration of the distal part of the planarian pharynx. Development Genes and Evolution, 2001. 211(1): p. 2–9.

26. Morita, M. and J.B. Best, Electron microscopic studies of planarian regeneration. IV. Cell division of neoblasts in *Dugesia dorotocephala*. Journal of Experimental Zoology, 1984. 229(3): p. 425–436.

27. Wenemoser, D. and P.W. Reddien, Planarian regeneration involves distinct stem cell responses to wounds and tissue absence. Dev Biol, 2010. 344(2): p. 979–91.

28. Xie, T. and A.C. Spradling, A Niche Maintaining Germ Line Stem Cells in the Drosophila Ovary. Science, 2000. 290(5490): p. 328–330.

29. Eisenhoffer, G.T., H. Kang, and A.S. Alvarado, Molecular Analysis of Stem Cells and Their Descendants during Cell Turnover and Regeneration in the Planarian Schmidtea mediterranea. Cell Stem Cell, 2008. 3(3): p. 327–339.

30. Scimone, M.L., et al., Two FGFRL-Wnt circuits organize the planarian anteroposterior axis. eLife, 2016. 5: p. e12845.

31. Wurtzel, O., I.M. Oderberg, and P.W. Reddien, Planarian Epidermal Stem Cells Respond to Positional Cues to Promote Cell-Type Diversity. Developmental Cell, 2017. 40(5): p. 491–504.e5.

32. Phelan, P., et al., Drosophila Shaking-B protein forms gap junctions in paired Xenopus oocytes. Nature, 1998. 391(6663): p. 181–184.

33. Oviedo, N.s.J. and M. Levin, smedinx-11 is a planarian stem cell gap junction gene required for regeneration and homeostasis. Development, 2007. 134(17): p. 3121–3131.

34. Nogi, T. and M. Levin, Characterization of innexin gene expression and functional roles of gap-junctional communication in planarian regeneration. Developmental Biology, 2005. 287(2): p. 314–335.

35. Lapan, Sylvain W. and Peter W. Reddien, Transcriptome Analysis of the Planarian Eye Identifies ovo as a Specific Regulator of Eye Regeneration. Cell Reports, 2012. 2(2): p. 294–307.

36. Lapan, S.W. and P.W. Reddien, dlx and sp6-9 Control Optic Cup Regeneration in a Prototypic Eye. PLoS Genetics, 2011. 7(8): p. e1002226.

37. Lei, K., et al., Pluripotency retention and exogenous mRNA introduction in planarian stem cells in culture. iScience, 2023. 26(2): p. 106001.

38. Cebrià, F. and P.A. Newmark, Planarian homologs of netrin and netrin receptor are required for proper regeneration of the central nervous system and the maintenance of nervous system architecture. Development, 2005. 132(16): p. 3691–3703.

39. Rozanski, A., et al., PlanMine 3.0—improvements to a mineable resource of flatworm biology and biodiversity. Nucleic Acids Research, 2018. 47(D1): p. D812–D820.

40. Untergasser, A., et al., Primer3--new capabilities and interfaces. Nucleic Acids Res, 2012. 40(15): p. e115.

41. Collins, J.J., 3rd, et al., Genome-wide analyses reveal a role for peptide hormones in planarian germline development. PLoS Biol, 2010. 8(10): p. e1000509.

42. Chandra, B., et al., Ets-1 transcription factor regulates glial cell regeneration and function in planarians. Development, 2023. 150(18).

43. King, R.S. and P.A. Newmark, In situ hybridization protocol for enhanced detection of gene expression in the planarian Schmidtea mediterranea. BMC Developmental Biology, 2013. 13(1): p. 8.

44. Ross, K.G., et al., Novel monoclonal antibodies to study tissue regeneration in planarians. BMC Developmental Biology, 2015. 15(1): p. 2.

45. Forsthoefel, D.J., F.A. Waters, and P.A. Newmark, Generation of cell type-specific monoclonal antibodies for the planarian and optimization of sample processing for immunolabeling. BMC Developmental Biology, 2014. 14(1): p. 45.

46. Jenkins, J.E. and R.H. Roberts-Galbraith, Heterotrimeric G proteins regulate planarian regeneration and behavior. Genetics, 2023. 223(4).

47. Schindelin, J., et al., Fiji: an open-source platform for biological-image analysis. Nature Methods, 2012. 9(7): p. 676–682.

48. Rouhana, L., et al., RNA interference by feeding in vitro–synthesized double-stranded RNA to planarians: Methodology and dynamics. Developmental Dynamics, 2013. 242(6): p. 718–730.

